# Log-linear scaling of TRPV4-KCNN4 transcripts tunes ROCK-dependent mechanotransduction in a DCIS progression model

**DOI:** 10.64898/2026.02.19.706850

**Authors:** Nathanael Ashby, Maxine Rubin, Robert G. Hawley, Inhee Chung

## Abstract

Mechanotransduction converts mechanical stress into cellular responses, yet how transcript abundance sets mechanotransduction capacity remains unclear. Using an isogenic MCF10A breast epithelial progression series in which crowding functionally inhibits plasma-membrane TRPV4 and triggers pro-invasive motility, we quantified how pathway mRNA levels relate to stress-evoked single-cell motility. TRPV4 (Ca²⁺-permeable mechanosensor) and KCNN4 (Ca²⁺-activated K⁺ channel) mRNA levels scaled log-linearly with motility under hyperosmotic stress or pharmacologic TRPV4 inhibition (both engaging the crowding-induced pro-invasive motility program) across a ∼600-fold TRPV4 mRNA range including isogenic and patient-derived DCIS lines (TRPV4: R²=0.89-0.92; KCNN4: R²=0.81-0.94). In contrast, bulk TRPV4 protein did not correlate with motility. Mechanistically, ROCK inhibition abolished stress-induced cortical actin-myosin organization and associated motility gains, identifying ROCK-dependent cortical contractility as a downstream effector. Notably, log-linear scaling was restricted to membrane channels and did not extend to tested cytosolic effectors, suggesting hierarchical transcript regulation may shape heterogeneous DCIS stress responsiveness in this model system.

## Introduction

Mechanotransduction pathways convert mechanical stimuli into cellular responses, yet how transcript abundance relates to functional output capacity remains poorly understood. This gap is especially relevant in ductal carcinoma in situ (DCIS), a non-invasive breast neoplasm with highly unpredictable clinical behavior^1–4^. DCIS cells proliferate within intact breast ducts^5–7^, where ductal confinement generates sustained crowding stress^8,9^. Although many DCIS lesions remain indolent, a subset progresses to invasive breast cancer^10^, and emerging evidence suggests that stress-adaptation capabilities may influence progression risk^8,11^. Defining the molecular basis of mechanotransduction capacity in DCIS could therefore reveal a mechanistic principle governing stress-driven invasive progression^12^.

We recently identified a TRPV4-centered mechanotransduction program in pro-invasive DCIS cell models using proteomic, biochemical, biophysical, and optical imaging approaches^8^. TRPV4 is a mechanosensitive, Ca²⁺-permeable ion channel that can contribute to basal Ca²⁺ influx^13,14^. In pro-invasive DCIS models, cell crowding functionally inhibits TRPV4 at the plasma membrane, reducing Ca²⁺ influx and initiating a stress-adaptation program marked by cell shrinkage, cortical stiffening, increased cortical actin and actin stress fiber formation, and compensatory TRPV4 relocalization to the plasma membrane^8^. These changes enhance single-cell motility and cell penetration capacity. Notably, both hyperosmotic stress and pharmacologic TRPV4 inhibition recapitulate this phenotype, driving TRPV4 plasma membrane relocalization, actin reorganization, and enhanced motility, thereby converging on the same mechanotransduction output as cell crowding^8^.

Using an isogenic MCF10A progression series^15,16^, we found that this stress-adaptation program is selectively engaged in pro-invasive DCIS (MCF10DCIS.com)^15^ and more mildly in invasive cancer (MCF10CA1a)^17^ cell lines, while notably diminished in less aggressive atypical ductal hyperplasia-like (MCF10AT1)^17^ and normal mammary epithelial (MCF10A) cells. This selectivity suggests that the response represents an acquired capability associated with invasive breast cancer progression^18^. Because hyperosmotic stress robustly engages this pathway, it provides a quantifiable readout of mechanotransduction capacity through single-cell motility measurements, which are technically impractical under crowding conditions^8^. Exploratory translational studies further suggest that pathology-dependent TRPV4 cellular distribution patterns may improve histopathologic classification when integrated with deep learning approaches^19^. Moreover, stress-induced plasma membrane relocalization of TRPV4, observed in patient DCIS specimens^8^, associates with progression to invasive disease^11^, though this relationship awaits comprehensive multi-institutional validation.

This mechanistic understanding revealed a paradox: while TRPV4 is essential for mechanotransduction as evidenced by partial knockdown reducing pathway output^8^, bulk TRPV4 protein abundance did not correlate with functional capacity across cell models^8^. This disconnect contrasts sharply with oncogene-driven cancers like HER2-amplified breast cancer, where protein overexpression directly predicts therapeutic vulnerability^20,21^. Why does total cellular TRPV4 protein fail to predict mechanotransduction capacity?

The answer likely lies in TRPV4’s dynamic trafficking behavior. Channel activation promotes internalization from the plasma membrane^8,22^, while channel inactivation during stress drives compensatory relocalization to the membrane^8^. Bulk protein abundance may vary with functional state, potentially limiting its predictive value. We reasoned that transcript abundance might provide a more stable readout: unlike protein levels, which vary with synthesis, trafficking, and degradation dynamics, mRNA abundance may better reflect the cell’s intrinsic capacity to deploy functional mechanosensor channels under stress.

Here we test whether TRPV4 mRNA abundance quantitatively predicts mechanotransduction output across the isogenic progression series and an extended panel including two patient-derived DCIS lines. We measure stress-evoked changes in single-cell motility under hyperosmotic conditions that engage the TRPV4-centered program and converge on the same output as cell crowding. If TRPV4 mRNA quantitatively predicts mechanotransduction output, the functional form of this relationship could reveal organizing principles governing mechanotransduction capacity. We address three questions: (i) Does TRPV4 mRNA correlate with functional output where bulk protein does not? (ii) What is the quantitative form of the transcript-output relationship? (iii) Is TRPV4 transcript abundance the dominant predictor, or do downstream effector transcripts augment prediction?

## Results

### TRPV4 mRNA exhibits log-linear scaling with mechanotransduction output

To test whether TRPV4 transcript abundance predicts mechanotransduction capacity, we measured stress-evoked single-cell motility changes compared to control conditions and TRPV4 mRNA across an isogenic MCF10A progression series representing normal epithelium (MCF10A), atypical ductal hyperplasia (MCF10AT1), high-grade DCIS (MCF10DCIS.com), and invasive carcinoma (MCF10CA1a), supplemented with two patient-derived DCIS lines (ETCC-06, ETCC-10)^23^ examined in our previous study^8^. Consistent with prior observations, stress-evoked motility responses were most pronounced for MCF10DCIS.com, with MCF10CA1a exhibiting a milder response. RNA-sequencing revealed that TRPV4 mRNA spanned >600-fold across the six-line panel, ranging from 154-fold lower in MCF10A and 622-fold lower in ETCC-06 relative to MCF10DCIS.com (**Figure 1A**).

**Figure 1.**
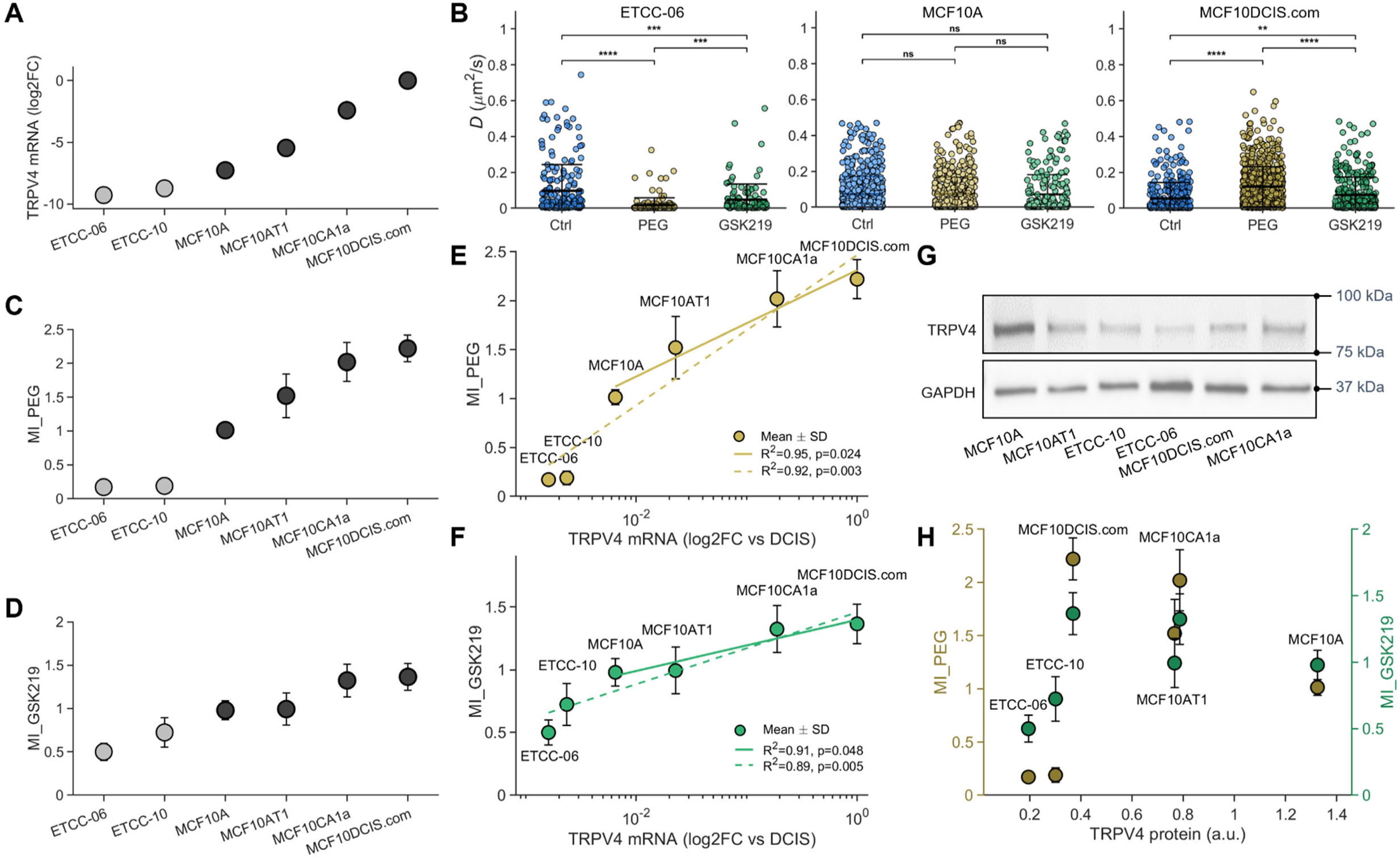
TRPV4 mRNA, but not TRPV4 protein, exhibits log-linear scaling with mechanotransduction output. **(A)** TRPV4 mRNA expression across a six-line breast cancer progression panel. Isogenic MCF10A series (black circles) and patient-derived DCIS lines (gray circles) shown as log2 fold-change (FC) relative to MCF10DCIS.com. TRPV4 mRNA spans a >600-fold range (∼622-fold across the six-line panel). Log2FC values (cell line/MCF10DCIS.com): ETCC-06, −9.28; ETCC-10, −9.04; MCF10A, −7.27; MCF10AT1, −5.23; MCF10CA1a, −2.18; MCF10DCIS.com, 0. **(B)** Representative single-cell diffusivity (*D*) distributions under control (ctrl, blue), hyperosmotic stress (PEG300, Δ74.4 mOsm/L, gold), and pharmacologic TRPV4 inhibition (GSK2193874, 1 nM, green) conditions. Three cell lines represent negatively responsive (ETCC-06), non-responsive (MCF10A), and highly responsive (MCF10DCIS.com) phenotypes. Statistical significance determined by two-sided Mann-Whitney U test: ns, not significant; *p<0.05; **p<0.01; ***p<0.001; ****p<0.0001. Each point is one cell; >100 cells total were tracked per condition across two independent experiments. **(C–D)** Motility indices (MI=<*D*_treatment>/<D_CONTROL>) for hyperosmotic stress (MI_PEG, **C**) and TRPV4 inhibition (MI_GSK219, **D**) across all six cell lines. Mean±SD; >100 cells total were tracked per condition across two independent experiments. **(E)** Log-linear relationship between TRPV4 mRNA and MI_PEG (gold circles). Isogenic MCF10A series (solid gold line, n=4) shows robust correlation (R²=0.95, p=0.024). Extended panel including patient-derived lines (dashed gold line, n=6) maintains significance (R²=0.92, p=0.003). **(F)** Log-linear relationship between TRPV4 mRNA and MI_GSK219 (green circles). Isogenic series (solid green line, n=4) shows correlation (R²=0.91, p=0.048), validating the mRNA–output relationship with an orthogonal trigger. Extended panel (dashed green line, n=6): R²=0.89, p=0.005. **(G)** TRPV4 protein expression by Western blot. Cells cultured at normal density to match motility assay conditions. GAPDH: loading control. **(H)** Bulk TRPV4 protein abundance (from densitometry of the WB data) versus mechanotransduction output for both triggers. Dual y-axes show MI_PEG (gold, left) and MI_GSK219 (green, right) as functions of normalized TRPV4 protein expression. No significant correlation observed (n=6; MI_PEG vs protein: R²=0.09, p=0.57; MI_GSK219 vs protein: R²=0.12, p=0.50).

We engaged the TRPV4-centered mechanotransduction program using two perturbations previously shown to engage the same pro-motility program as cell crowding while remaining compatible with single-cell tracking at normal density: hyperosmotic stress (PEG300; Δ74.4 mOsm/L) and pharmacologic TRPV4 inhibition (GSK2193874, 1 nM)^8^. Both triggers can be applied at normal cell density, enabling tracking of individual cell movements, which is technically impractical under crowding conditions. Single-cell motility was quantified by time-lapse confocal microscopy to obtain diffusivity coefficients (*D*) under control and stress conditions, using a linear mean-squared displacement (MSD) fit (**Methods**, **Figure S1**). Single-cell diffusivity (*D*) scatter plots for three cell lines spanning the phenotypic spectrum are shown in **Figure 1B**, with full plots for all six lines in **Figure S2**.

To quantify stress-evoked motility changes, we calculated motility indices (MI) for each line as the ratio of mean single-cell diffusivity under treatment versus control (MI_PEG=<*D*_PEG>/<*D*_control>; MI_GSK219=<*D*_GSK219>/<*D*_control>). Across the panel, MI_PEG ranged from 0.17 (ETCC-06) to 2.22 (MCF10DCIS.com), and MI_GSK219 ranged from 0.50 (ETCC-06) to 1.37 (MCF10DCIS.com) (**Figure 1C, D**). Lines exhibited graded responses spanning a continuous spectrum: the strongest responders increased motility under both triggers (MCF10DCIS.com: MI_PEG=2.22±0.20, MI_GSK219=1.37±0.16; MCF10CA1a: MI_PEG=2.01±0.29, MI_GSK219=1.32±0.19), intermediate responders showed modest or no changes (MCF10AT1: MI_PEG=1.52±0.32, MI_GSK219=0.99±0.19; MCF10A: MI_PEG=1.01±0.07, MI_GSK219=0.97±0.11), and negative-responders exhibited reduced motility under PEG300 (ETCC-10: MI_PEG=0.19±0.07; ETCC-06: MI_PEG=0.17±0.04) and TRPV4 inhibition (ETCC-10: MI_GSK219=0.72±0.17; ETCC-06: MI_GSK219=0.50±0.10).

We next examined the quantitative relationship between TRPV4 mRNA and mechanotransduction output (MI). Across the isogenic series, TRPV4 mRNA exhibited log-linear scaling with MI_PEG (R²=0.95, p=0.024, n=4) (**Figure 1E**, solid gold line), in which fold-changes in mechanosensor transcript correspond to proportional changes in output. An orthogonal trigger produced the same relationship: TRPV4 mRNA also scaled log-linearly with MI_GSK219 (R²=0.91, p=0.048, n=4) (**Figure 1F**, solid green line). Alternative model fits (linear or log-log) showed lower R² values for both the isogenic and extended series (**Figure S3**). The concordance between hyperosmotic stress and direct channel inhibition, two mechanistically distinct triggers that converge on the mechanotransduction output, supports TRPV4 transcript abundance as a robust predictor of mechanotransduction capacity.

We then tested whether the patient-derived DCIS lines examined in our previous study^8^ follow this relationship. Including ETCC-06 and ETCC-10 extended the analysis across the full six-line panel (**Figure 1E**, **F**; dashed lines). For MI_PEG, the extended panel maintained strong correlation (R²=0.92, p=0.003, n=6). For MI_GSK219, the extended panel similarly showed strong correlation (R²=0.89, p=0.005, n=6), maintaining the log-linear relationship across both triggers.

### Bulk TRPV4 protein does not predict mechanotransduction output

In contrast to the strong mRNA-output relationship, bulk TRPV4 protein abundance measured by Western blot did not correlate with mechanotransduction capacity, consistent with our previous study^8^. TRPV4 protein levels varied only modestly across cell lines (**Figure 1G**, **Supplemental Figure S4**) and showed no relationship with output for either trigger (**Figure 1H**; MI_PEG: R²=0.09, p=0.57; MI_GSK219: R²=0.12, p=0.50; n=6). Together, these results indicate that TRPV4 mRNA, not bulk protein, quantitatively scales with mechanotransduction capacity.

The strength of the TRPV4 mRNA-output relationship, together with the lack of predictive power for bulk TRPV4 protein, motivated testing whether transcript-level scaling is concentrated at the plasma membrane sensor layer or extends to downstream cytoplasmic effectors. We therefore first established which downstream pathways are required for execution of the mechanotransduction phenotype. Because crowding induces cortical and stress-fiber remodeling in pro-invasive DCIS cells^8^, we tested whether Rho/ROCK signaling^24,25^ is functionally required for stress-evoked actin organization and motility increase.

### ROCK-dependent cortical contractility mediates stress-evoked motility

We used ROCK inhibition (Y-27632, 50 µM) to define which aspects of stress-evoked actin remodeling require ROCK activity. Using our in-house custom-built 3D-structured illumination microscopy (3D-SIM)^26^, we observed that hyperosmotic stress (PEG300) produced minimal changes in ventral stress fibers (**Figure 2A**), in contrast to the prominent stress fibers observed under cell-crowding stress^8^. This distinction suggests that stress fiber assembly during crowding may additionally depend on cell-cell contact signaling. Instead, we observed increased cortical actin reorganization under PEG300 stress and therefore examined ROCK’s role in cortical contractility.

**Figure 2.**
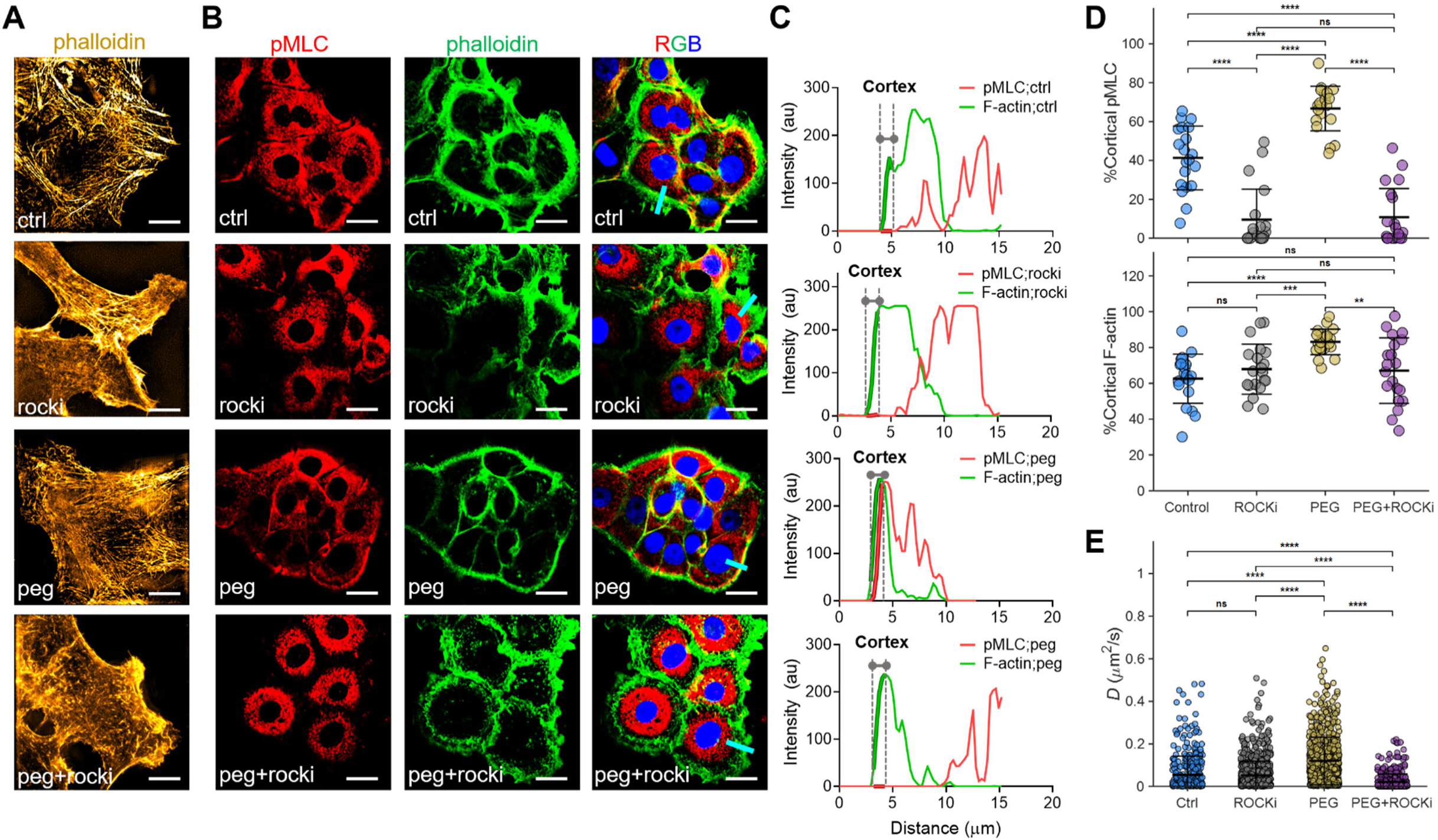
ROCK-dependent cortical actin reorganization mediates mechanotransduction-induced motility enhancement. **(A)** Representative 3D-SIM images of F-actin (phalloidin, yellow) in MCF10DCIS.com cells under control conditions (ctrl), ROCK inhibition (Y-27632, 50 μM; ROCKi), hyperosmotic stress (PEG300 Δ74.4 mOsm/L; PEG), and combined treatment (PEG+ROCKi). Scale bars: 10 μm. **(B)** Confocal images of phosphorylated myosin light chain (pMLC, red) localization showing ROCK-dependent myosin activation at the cell cortex under hyperosmotic stress. ROCK inhibition abolishes pMLC cortical enrichment. Merge panels show F-actin (green), pMLC (red), and DAPI (blue). Scale bars: 20 μm. **(C)** Phalloidin staining (green) with corresponding intensity line scans across cell diameter (white arrows) for control and treated conditions. Merge panels show F-actin (green), pMLC (red), and nuclear DNA (RGB, blue). Hyperosmotic stress (PEG) induces cortical F-actin and pMLC enrichment, which is abolished by ROCK inhibition (PEG+ROCKi). Distance measured from cell edge; cortex defined as 1.5 μm from periphery. Scale bars: 20 μm. **(D)** Quantification of cortical pMLC intensity (top) and cortical F-actin content (bottom). Hyperosmotic stress induces cortical enrichment: cortical pMLC increased from 41.3±16.4% (Control) to 66.7±11.5% (PEG, p<0.0001), and cortical F-actin increased from 62.6±13.7% (Control) to 83.1±7.1% (PEG, p<0.0001). ROCK inhibition prevents stress-induced enrichment: PEG+ROCKi pMLC 10.8±14.8% (p<0.0001 vs PEG) and PEG+ROCKi F-actin 67.1±18.3% (p<0.0001 vs PEG, ns vs Control). ROCK inhibition alone reduces basal cortical pMLC to 9.5±15.6% (p<0.0001 vs Control) but does not affect cortical F-actin (67.9±14.0%, ns vs Control). Each point: single cell; n≥30 cells per condition from 3 independent experiments. Horizontal bars: mean ± SD. *p<0.05, **p<0.01, ****p<0.0001; ns, not significant (Mann-Whitney U test for pairwise comparisons). **(E)** Single-cell motility diffusion coefficient (*D*) measurements demonstrate that ROCK inhibition abolishes mechanotransduction-induced motility enhancement. Hyperosmotic stress increases motility (Control: 0.054±0.088 µm²/s; PEG: 0.120±0.111 µm²/s, p<0.0001), which is completely blocked by ROCK inhibition (PEG+ROCKi: 0.021±0.037 µm²/s, p<0.0001 vs PEG, ns vs Control). ROCK inhibition alone has minimal effect (ROCKi: 0.052±0.066 µm²/s, ns vs Control). Each point: single cell; ≥100 cells total were tracked per condition across two independent experiments. *p<0.05, **p<0.01, ****p<0.0001; ns, not significant (Mann-Whitney U test for pairwise comparisons).

We assessed cortical contractility by examining phospho-myosin light chain 2 (pMLC) localization relative to F-actin^27^. Under control conditions, pMLC showed moderate cortical distribution with partial overlap with cortical F-actin (**Figure 2B**, row 1), whereas ROCK inhibition redistributed pMLC to the cytoplasm (row 2). PEG300 induced striking cortical pMLC enrichment that tightly colocalized with cortical F-actin, which also became enriched at the cell periphery, forming a circumferential contractile ring (row 3). Line profiles confirmed edge-localized enrichment, with pMLC peaks positioned just interior to F-actin peaks (∼ 0.5 µm inward), consistent with myosin motors acting on a cortical actin scaffold (**Figure 2C**). ROCK inhibition abolished PEG300-induced cortical pMLC enrichment (row 4), eliminating the cortical peaks.

We quantified both cortical pMLC and cortical F-actin as the fraction of total cellular signal within a ∼1.5 µm peripheral band defined by phalloidin staining (**Figure 2D**; cortex masks shown in **Figure 2C**). PEG300 increased cortical pMLC from 41.3±16.4% to 66.7±11.5% (p<0.0001). ROCK inhibition reduced basal cortical pMLC (9.5±15.6%, p<0.0001 vs control) and also prevented PEG300-induced enrichment (PEG+ROCKi: 10.8±14.8%, p<0.0001 vs PEG; ns vs ROCKi alone). Similarly, PEG300 also increased cortical F-actin from 62.6±13.7% to 83.1±7.1% (p<0.0001 vs control), while ROCK inhibition blocked this enrichment (PEG+ROCKi: 67.1±18.3%, p<0.0001 vs PEG; ns vs control). ROCK inhibition alone did not alter baseline cortical F-actin distribution (67.9±14.0%, ns vs control). ROCK activity is therefore required for both basal cortical pMLC localization and stress-induced cortical enrichment.

Consistent with these PEG300 findings, pharmacologic TRPV4 inhibition (GSK2193874, 10 nM for imaging to enhance signal consistency; motility assays used 1 nM throughout) also promoted cortical enrichment of pMLC and F-actin **(Supplemental Figure S5**). Quantification as in **Figure 2D** showed that GSK219 increased cortical pMLC from 41.3±16.4% (control) to 55.9±16.9% (p=0.040). ROCK inhibition reduced pMLC enrichment and attenuated the GSK219 response (GSK219+ROCKi: 24.7±14.6%; p=0.003 vs GSK219). Similarly, GSK219 increased cortical F-actin from 62.6±13.7% (control) to 76.8±12.2% (p=0.012), while ROCK inhibition reduced this enrichment (GSK219+ROCKi: 61.4±11.3%, p=0.017 vs GSK219). Together, these results indicate that TRPV4 inhibition engages ROCK-dependent cortical myosin activation and promotes cortical actin enrichment, paralleling the PEG300 response.

To test the functional necessity, we measured single-cell diffusivity (**Figure 2E**). PEG300 increased motility (0.120±0.111 µm²/s) compared to control (0.054±0.088 µm²/s, p<0.0001). ROCK inhibition alone did not alter baseline motility (0.052±0.066 µm²/s, p=0.65 vs control), but it abolished PEG300-evoked enhancement (PEG+ROCKi: 0.021±0.037 µm²/s, p<0.0001 vs PEG). Thus, ROCK-dependent cortical contractility, manifest as coordinated enrichment of both F-actin and pMLC at the cell cortex, is required for stress-evoked motility.

### Cytoplasmic effector transcripts do not predict capacity despite functional requirement

Having established ROCK as essential for execution, we sought to examine if cytoplasmic effector transcript abundance also predicts mechanotransduction capacity. ROCK2 mRNA levels varied minimally across the panel (2.8-fold range) and did not correlate with mechanotransduction output for either trigger (MI_PEG: R²=0.51, p=0.174; MI_GSK219: R²=0.40, p=0.254; n=5; **Supplemental Figure S6**; **Supplemental Table S1**). These results support a two-tier organization in which mechanotransduction capacity is determined by mechanosensor transcript abundance (TRPV4: 622-fold range, R²>0.89, p<0.005; **Figure 1E, F**), while functionally required downstream effectors such as ROCK2 show limited transcript variation and do not explain between-line differences in output (2.8-fold range, R²<0.52, p>0.17).

### TRPV4 and KCNN4 form a co-regulated mechanosensor module at the plasma membrane

Because mechanotransduction capacity tracked TRPV4 transcript abundance but not bulk TRPV4 protein or ROCK2 mRNA, we asked whether capacity is selectively encoded at the level of plasma-membrane sensors, or whether other pathway components also exhibit transcript-output scaling. To address this, we surveyed candidate genes spanning the mechanotransduction axis: mechanosensitive ion channels (TRPV4, KCNN4, PIEZO1, PIEZO2)^8^, cytoplasmic Rho/ROCK signaling components (RHOB, RHOC, ROCK2)^28^, and mechanosensitive transcriptional regulators (YAP1, WWTR1/TAZ)^29,30^. We selected KCNN4 based on our prior proteomics dataset^8^ and its function as a Ca²⁺-activated K⁺ channel positioned to couple TRPV4-mediated Ca²⁺ influx to downstream signaling (**Figure 3A** shows expression across the panel; **Figure 3B** summarizes expression range and correlation strengths; correlations with MI are shown in **Figure 3C–D**; full statistics are provided in **Supplemental Table S1**). Across the six-line panel, PIEZO1/2 were absent from the vendor DE result exports used to extract log2 fold-change values and therefore analysis was not available (missing values do not imply zero expression in the reference). Among detected transcripts, KCNN4 spanned an 86-fold range across all six lines, whereas TRPV4 spanned 622-fold.

**Figure 3.**
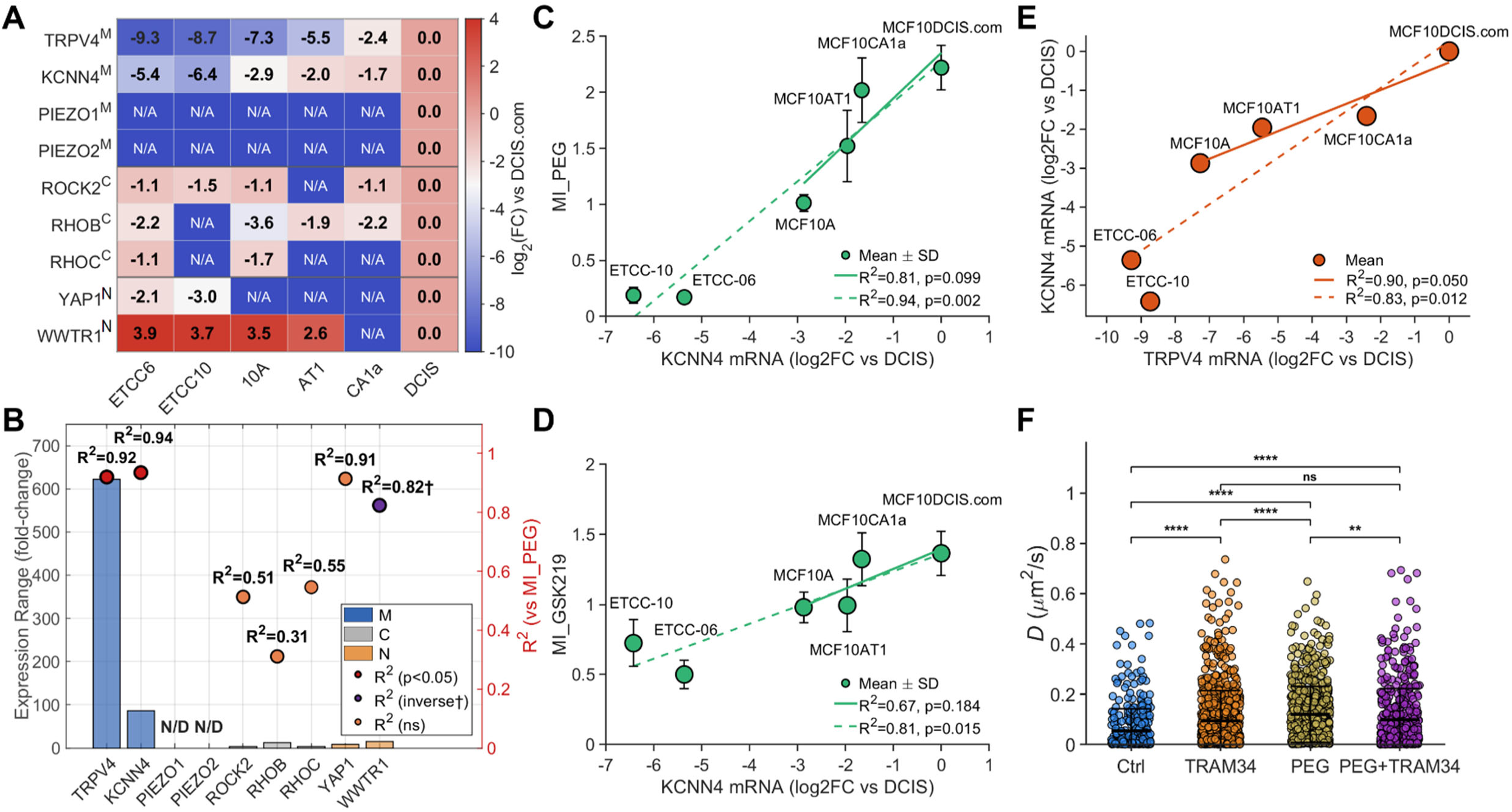
TRPV4 and KCNN4 form a transcriptionally co-regulated mechanosensor module. **(A)** Heatmap of candidate mechanotransduction genes across six cell lines ordered by increasing mechanotransduction capacity (ETCC-06 lowest to MCF10DCIS.com highest). Values are log2(mRNA_cell line/mRNA_MCF10DCIS.com). NA indicates log2 fold-change values that were not available from the differential-expression outputs used for fold-change extraction, resulting in gene-specific n<6 for some genes (e.g., PIEZO1/2). Superscripts indicate predominant localization: M, membrane; C, cytosolic; N, nuclear. **(B)** Summary correlations between candidate gene expression (log2FC vs MCF10DCIS.com) and mechanotransduction output (MI_PEG). Bars denote expression dynamic range across the panel; overlaid points report R² from linear regression of gene expression versus MI_PEG (red, p<0.05; orange, p≥0.05; purple, significant inverse correlation (p<0.05)). Among surveyed candidates, TRPV4 (R²=0.92, p=0.003) and KCNN4 (R²=0.94, p=0.002) show the strongest positive predictive relationships. **(C)** KCNN4 mRNA abundance predicts MI_PEG with log-linear scaling. Points represent individual cell lines; error bars: mean±SD. Dashed line: full panel regression (n=6, R²=0.94, p=0.002); solid line: isogenic MCF10A series (n=4, R²=0.81, p=0.099). **(D)** KCNN4 mRNA abundance predicts MI_GSK219 with similar log-linear scaling. Same plotting conventions as in **(C)**. Dashed line: full panel (n=6, R²=0.81, p=0.015); solid line: isogenic series (n=4, R²=0.67, p=0.184). **(E)** TRPV4 and KCNN4 mRNA levels are strongly correlated across the cell-line panel, indicating coordinated transcriptional regulation. Dashed line: full panel (n=6, R²=0.83, p=0.012); solid line: isogenic MCF10A series (n=4, R²=0.90, p=0.050). **(F)** Functional validation: pharmacologic KCNN4 inhibition (TRAM-34, 5 μM) phenocopies the motility increase induced by hyperosmotic stress (PEG300; Δ74.4 mOsm/L). Distributions show single-cell diffusion coefficients (*D*) for control, TRAM-34, PEG, and PEG+TRAM-34. TRAM-34 (0.094±0.120 µm²/s) and PEG (0.120±0.111 µm²/s) each increased motility versus control (0.054±0.088 µm²/s; both p<0.0001), with no additive effect in the combined condition (PEG+TRAM-34: 0.099±0.124 µm²/s, ns vs either alone). Each point is one cell; >100 cells total were tracked per condition across two independent experiments. Statistics: Mann-Whitney U test. **p<0.01; ****p<0.0001; ns, not significant.

We then tested correlations between transcript abundance and mechanotransduction output (MI_PEG and MI_GSK219) across the panel using linear regression (p-values test a nonzero regression slope). Because log2FC values were unavailable for some genes in one or more lines, n reflects the number of lines with available values. Beyond TRPV4 (MI_PEG: R²=0.92, p=0.003; MI_GSK219: R²=0.89, p=0.005; n=6), only KCNN4 showed significant correlation with both triggers (MI_PEG: R²=0.94, p=0.002; MI_GSK219: R²=0.81, p=0.01; n=6) (**Figure 3C**, **D**). Like TRPV4, KCNN4 transcript levels showed log-linear scaling with output, albeit over a shorter expression range than TRPV4 (86-fold vs 622-fold). In contrast, Rho GTPase signaling components showed no significant correlations: ROCK2 (MI_PEG: R²=0.51, p=0.10; MI_GSK219: R²=0.40, p=0.25; n=5), RHOB (MI_PEG: R²=0.31, p=0.33; MI_GSK219: R²=0.19, p=0.46; n=5), and RHOC (MI_PEG: R²=0.55, p=0.47; MI_GSK219: R²=0.38, p=0.58; n=3; **Supplemental Table S1**). Similarly, mechanosensitive transcriptional regulators showed no positive scaling with capacity, including YAP1 (MI_PEG: R²=0.91, p=0.189; MI_GSK219: R²=0.74, p=0.345; n=3), and WWTR1/TAZ, which exhibited an inverse relationship.

Because TRPV4 is a Ca²⁺-permeable mechanosensor and KCNN4 encodes a Ca²⁺-activated K⁺ channel, we next asked whether these channels are coordinately regulated and functionally coupled. Consistent with their shared scaling with MI (**Figure 3C, D**), KCNN4 also tracked TRPV4 expression across cell lines, suggesting a shared regulatory program (**Figure 3E**). Indeed, TRPV4 and KCNN4 mRNA levels were strongly correlated across the panel (R²=0.83, p=0.012), indicating coordinated transcriptional regulation (**Figure 3E**). To test whether KCNN4 participates in the same mechanotransduction pathway, we inhibited KCNN4 with TRAM-34 (5 μM). KCNN4 inhibition increased motility to levels indistinguishable from hyperosmotic stress (Control: 0.054±0.088 µm²/s; TRAM-34: 0.094±0.120 µm²/s; PEG300: 0.120±0.111 µm²/s; both p<0.0001 vs Control, ns between TRAM-34 and PEG300), and combining treatments produced no additive effect (PEG300+TRAM-34: 0.099±0.124 µm²/s, ns vs either alone) (**Figure 3F**). This non-additivity supports a model in which TRPV4 and KCNN4 operate within the same mechanotransduction pathway rather than contributing independent, additive inputs. Furthermore, cytoskeletal analysis also showed that TRAM-34 induces cortical actin reorganization similar to PEG300 (**Supplemental Figure S7**), consistent with engagement of the same cortical remodeling phenotype defined under PEG stress. Together, these results support a model in which TRPV4 and KCNN4 form a coordinately regulated plasma-membrane ion-channel module: both channels scale log-linearly with mechanotransduction capacity at the transcript level, and inhibition of either channel phenocopies hyperosmotic stress, triggering the same pro-invasive program of cortical contractility and enhanced motility. Consistent with this coupling, KCNN4 provided no additional predictive value beyond TRPV4 in multiple regression models (p>0.09 for both triggers; **Supplemental Table S2**). Together, these findings support a two-tier organizational framework for mechanotransduction capacity in DCIS (**Figure 4**).

**Figure 4.**
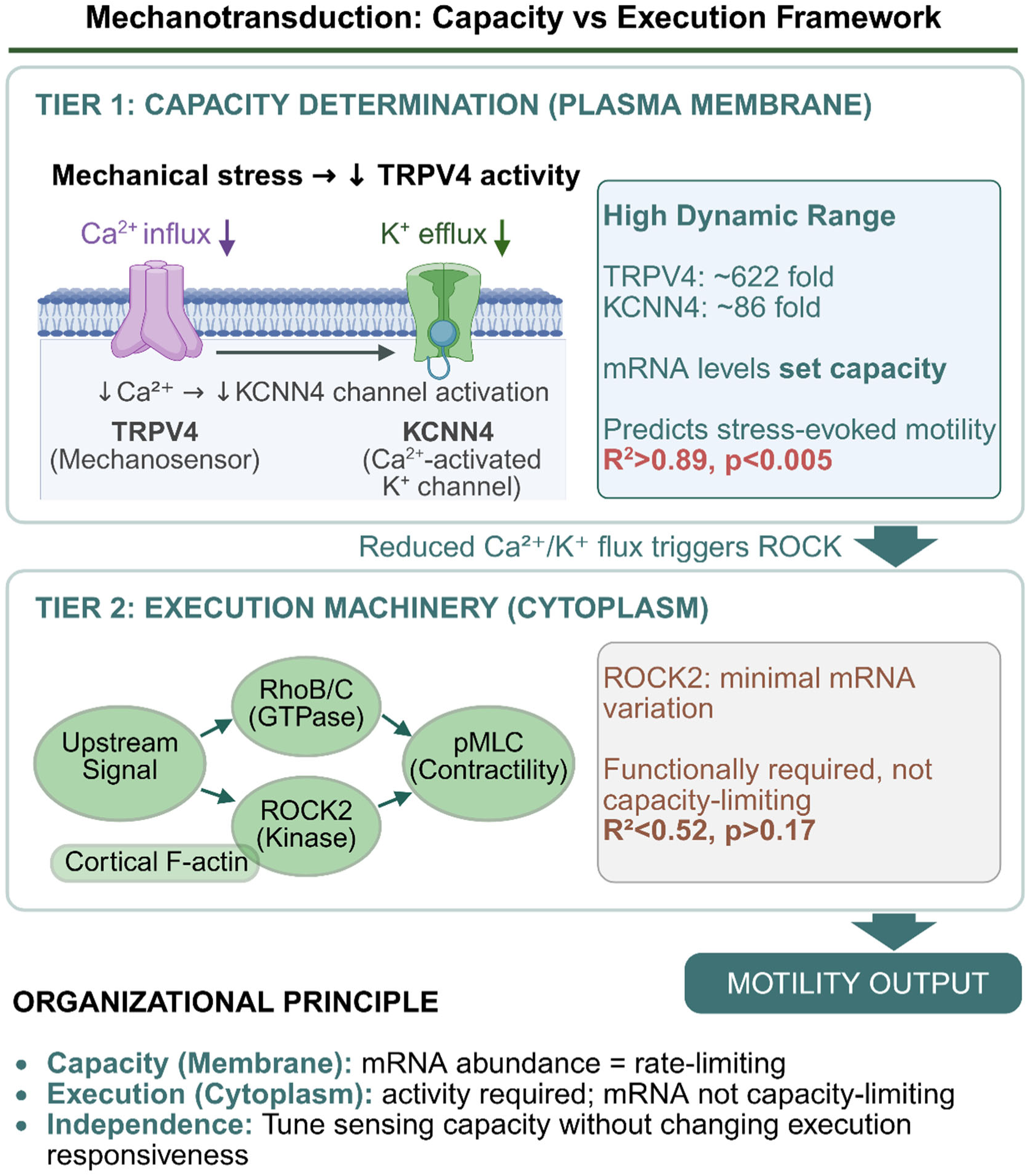
Two-tier architecture of mechanotransduction capacity in DCIS. Tier 1 (plasma membrane): TRPV4 and KCNN4 transcript abundance spans ∼86–622-fold across the progression panel and predicts stress-evoked motility with log-linear scaling (R²>0.89, p<0.005). Mechanical stress (and pharmacologic TRPV4 inhibition) reduces TRPV4-mediated Ca²⁺ entry and functionally attenuates the TRPV4–KCNN4 module, yielding a reduced Ca²⁺/K⁺ flux signal that triggers ROCK-dependent cortical contractility. Tier 2 (cytoplasm): ROCK2 and Rho GTPase pathway effector transcripts vary minimally and do not predict motility output (R²<0.52, p>0.17) despite functional requirement. Together, mechanotransduction capacity is encoded at the membrane sensor level, whereas downstream execution machinery is broadly available and regulated primarily at the activity level.

## Discussion

DCIS progression risk remains difficult to predict because conventional proliferation or invasion markers do not reliably identify the subset that becomes pro-invasive under ductal confinement^31,32^. In our prior work^8^, we showed that DCIS models differ in their response to mechanical stress, where some engage a TRPV4-dependent program that promotes motility, whereas others do not. Although TRPV4 was required for this response, bulk TRPV4 protein abundance did not track mechanotransduction capacity, raising a central question: why is TRPV4 essential, yet total protein not predictive?

Here we resolve this discrepancy by identifying a quantitative link between transcript abundance and stress-evoked behavioral output. Across a DCIS-focused progression panel, TRPV4 mRNA spanned >600-fold and predicted mechanotransduction capacity, scaling log-linearly with single-cell motility under orthogonal perturbations (hyperosmotic stress and pharmacologic TRPV4 inhibition; R²>0.90). In contrast, bulk TRPV4 protein showed no correlation. Although the panel is limited in size, the relationship is reproduced with two orthogonal perturbations and persists when patient-derived DCIS lines extend the dynamic range. This log-linear relationship, where fold-changes in transcript correspond to proportional changes in functional output, is consistent with Weber-Fechner-like scaling observed in other biological sensing systems^33,34^. In sensory systems, Weber-Fechner-like scaling reflects logarithmic compression, in which equal stimulus ratios produce equal response increments, enabling wide dynamic range while preserving sensitivity to relative change. Our findings suggest that a similar principle can apply to mechanotransduction capacity: TRPV4-KCNN4 transcript abundance functions as a quantitative gain-setting layer, such that fold-changes in mechanosensor transcripts map to proportional increments in stress-evoked motility output. This framework is conceptually distinct from classic thresholded or saturating signaling behaviors^35,36^ and provides a parsimonious explanation for graded, heterogeneous stress responsiveness across DCIS progression states.

Mechanistically, both hyperosmotic stress and pharmacological TRPV4 inhibition, which mimics the functional TRPV4 inactivation at the plasma membrane that occurs during cell crowding^8^, converge on ROCK-dependent cortical contractility. Both triggers induced cortical enrichment of pMLC and F-actin, and ROCK inhibition suppressed this cortical remodeling; in motility assays, ROCK inhibition abolished the PEG300-evoked motility increase, identifying ROCK-dependent cortical contractility as a shared execution mechanism.

Pathway-wide transcript analysis further supports a two-tier organization. Beyond TRPV4, KCNN4 mRNA showed significant predictive power with a log-linear relationship with stress induced motility increases (R²>0.8), and TRPV4 and KCNN4 transcripts were strongly correlated across cell lines (R²=0.83). KCNN4 was selected based on prior proteomics^8^ and its function as a Ca²⁺-activated K⁺ channel positioned to couple TRPV4-mediated Ca²⁺ influx to downstream signaling. Functional coupling was supported by pharmacologic inhibition: KCNN4 blockade phenocopied stress-evoked motility and cortical remodeling with no additivity, consistent with TRPV4, KCNN4, and hyperosmotic stress operating within the same pathway. This coordinated TRPV4-KCNN4 module parallels findings in glioma invasion, where Ca²⁺-permeable channels and Ca²⁺-activated K⁺ channels (including KCNN4) coordinate regulated volume changes required for migration through confined spaces^37,38^, providing a mechanistic precedent for ion-channel-mediated volume regulation engaging downstream contractility pathways. In contrast, cytoplasmic effectors such as ROCK2 showed no transcript-output correlation despite functional requirement, indicating that capacity-associated variation is concentrated at the membrane sensor level, while downstream machinery is broadly available and engaged through post-translational activation.

The disconnect between transcript-level prediction and bulk protein abundance suggests that total protein is not a reliable proxy for the signaling-competent pool, consistent with regulation by trafficking, localization, and/or complex assembly^8^. Given the limited availability of patient-derived DCIS models for systematic mechanotransduction phenotyping, we focused on well-characterized lines enabling single-cell tracking and high-resolution cytoskeletal readouts. Future work should test whether this transcript-output relationship generalizes to larger patient-derived collections and whether spatially resolved measures of TRPV4/KCNN4 expression and/or subcellular TRPV4 localization improve prediction of progression-associated phenotypes^11,19^.

In sum, mechanotransduction capacity is encoded at the transcript level by a plasma membrane ion-channel module (TRPV4-KCNN4), while ROCK-dependent cortical contractility is essential for execution but not capacity-limiting at the transcript level. This two-tier architecture provides a framework for DCIS heterogeneity in stress responsiveness and helps explain why transcript-based measurements can outperform bulk protein assays in mechanosensing contexts.

## Data availability

RNA-seq data will be deposited in the NCBI Gene Expression Omnibus (GEO) upon acceptance. All other data supporting the findings are available in the supplemental information or from the corresponding author (I.C.) upon reasonable request.

## Acknowledgments

I.C. is supported by the George Washington Cancer Center and the Katzen Cancer Research Pilot Award. We acknowledge K. Yun for RNA extraction, and C. Chan, J. Kim, and A. Baker for their efforts in cell management.

## Author Contributions

I.C. conceived and supervised the project, secured funding, designed experiments, methods, and analyses, and wrote the manuscript with comments from all authors. N.A. conducted immunofluorescence imaging experiments, performed line profile analysis, and acquired single-cell motility movies. M.R. performed Western blot experiments and acquired single-cell motility movies. R.H. consulted on RNA-seq data analysis.

## Corresponding authors

Correspondence to Inhee Chung.

## Methods

### Cell Lines

MCF10A cells were purchased from ATCC, and MCF10AT1, MCF10DCIS.com, and MCF10CA1a cells were obtained from the Barbara Ann Karmanos Cancer Institute through a material transfer agreement. ETCC-06 and ETCC-10 cells were purchased from the Leibniz Institute DSMZ. All cell lines used in this study are of female origin. All cell lines were authenticated by the supplier, used at a low passage number (<15), and confirmed to be mycoplasma-free by PCR analysis. MCF10A and MCF10AT1 cells were maintained in DMEM/F12 medium supplemented with 5% horse serum, 20 ng/mL EGF, 0.5 μg/mL hydrocortisone, 10 μg/mL insulin, and 1% penicillin and streptomycin. Additionally, MCF10A cells were supplemented with 100 ng/mL cholera toxin. MCF10DCIS.com and MCF10CA1a cells were cultured in DMEM/F12 with 5% horse serum and 1% penicillin and streptomycin. ETCC-06 and ETCC-10 cells were maintained in RPMI 1640 supplemented with 10% fetal bovine serum (FBS) and 1% penicillin and streptomycin. All cells were cultured at 37°C in 5% CO₂.

### RNA Sequencing and Analysis

Total RNA was isolated from cultured cells using the RNAqueous kit (Thermo Fisher, AM1912) following the manufacturer’s protocol. RNA concentration and purity were assessed by NanoDrop spectrophotometry. RNA-seq library preparation and sequencing were performed by Azenta Life Sciences using a poly(A)-selected mRNA workflow. Libraries were sequenced on an Illumina HiSeq platform generating paired-end 2×150 bp reads. Raw reads were quality-filtered and adapter-trimmed using Trimmomatic v0.36, aligned to the human GRCh38 reference genome (Ensembl annotation) using the splice-aware aligner STAR v2.5.2b, and gene-level counts were quantified from uniquely mapped reads using featureCounts (Subread v1.5.2). Only unique reads mapping to exon regions were counted. Differential expression analysis was performed by Azenta using DESeq2, with each cell line compared against MCF10DCIS.com as a common reference. Statistical testing used the Wald test to generate p-values and log2 fold changes, with Benjamini–Hochberg correction for multiple testing. Genes with adjusted p-value <0.05 and absolute log2 fold change >1 were considered differentially expressed. For correlation analyses, log2 fold-change values (relative to MCF10DCIS.com) for candidate mechanotransduction genes were obtained from the Azenta differential-expression outputs and correlated with motility indices using linear regression as described in Results. When a candidate gene was not present in the differential-expression output for a given contrast, its log2 fold-change value was treated as missing and that gene/contrast datapoint was excluded from the corresponding regression; thus, the effective n could vary by gene and condition.

### Western Blot Analysis

Cells were lysed in cytoskeleton buffer (10 mM Tris pH 7.4, 100 mM NaCl, 1 mM EDTA, 1 mM EGTA, 1% Triton X-100, 10% glycerol, 0.1% SDS, 0.5% deoxycholate). Lysates were then mixed with SDS sample buffer (Bio-Rad) supplemented with reducing reagent and boiled at 100°C for 10 min. Proteins were separated on 4-15% SDS-PAGE gels and transferred to nitrocellulose membranes (Bio-Rad). Membranes were blocked in TBST (TBS + 0.1% Tween-20) containing 5% non-fat milk and incubated overnight at 4°C with primary antibodies: rabbit anti-TRPV4 (Proteintech, 84091-4-RR, 1:500) and mouse anti-GAPDH (Santa Cruz, sc-32233, 1:5,000). After three TBST washes (5 min each), membranes were incubated with HRP-conjugated goat anti-rabbit (Thermo Fisher Scientific 31460, 1:5,000) or goat anti-mouse (Thermo Fisher Scientific 31430, 1:5,000) secondary antibodies for 30 min at room temperature. Blots were visualized using Pico Plus chemiluminescent substrate (Thermo Fisher Scientific, 34580).

### Immunofluorescence

Cells were fixed with 4% paraformaldehyde (Fisher Scientific) for 20 min, followed by permeabilization with 0.1% saponin (Fisher Scientific) for 5 min at room temperature. After permeabilization, cells were washed three times in PBS and blocked in 0.1% saponin + 10% BSA in PBS at room temperature for 1 hr. Cells were incubated with primary antibodies overnight at 4°C in 0.1% saponin in PBS. The primary antibodies used were anti-Phospho-Myosin Light Chain 2 (Ser19) (pMLC, Cell Signaling 3671; lot 7; 1:200 dilution) and anti-TRPV4 (Abcam ab39260; 1:500 dilution). Cells were washed three times in PBS, incubated with fluorescent-tagged anti-rabbit Alexa Fluor 647 secondary antibodies (Thermo Fisher Scientific, A-31573, 1:3000) for 1 hr at room temperature, followed by ActinGreen 488 ReadyProbes Reagent (Thermo Fisher R37110; 1 drop per 1 mL PBS), and 300 nM DAPI (Thermo Fisher D3571) in PBS. Samples were then imaged by confocal microscopy.

### Confocal Imaging and 3D-SIM Super-Resolution Imaging

Spinning-disk confocal images were acquired using a Yokogawa spinning-disk confocal system (Andor Technology) installed on a Nikon Eclipse TE2000 inverted microscope with a 60×/1.49 NA Plan Apo objective (Nikon). Samples were illuminated using 488, 561, and 647 nm solid-state lasers (Andor Technology) and images were acquired with an iXon back-illuminated EMCCD camera (Andor Technology). Three-dimensional structured illumination microscopy (3D-SIM) was performed using our custom-built multi-angle-crossing structured illumination microscope (MAxSIM) without the height-controlled mirror^26^. The MAxSIM system that enables any dimensional SIM performances was built on a Zeiss Axio Observer inverted microscope platform with an ASI motorized stage and a 100×/1.46 NA oil immersion objective (Alpha Plan-APO, Zeiss). Two-color sequential imaging was performed using 488 nm and 647 nm excitation. Excitation was controlled by a software-controlled filter wheel (Finger Lakes Instrumentation) and grating patterns were generated using a spatial light modulator (Forth Dimension Display). Fluorescence images were collected using an sCMOS camera (Hamamatsu Flash 4.0) with 100 ms exposure time. 3D-reconstruction was performed using published custom software^39^. System control was performed using VSIM open-source software (HHMI). All images were processed using ImageJ (NIH).

### Cell Motility Assay

Cells were seeded in MatTek 35 mm dishes with No. 1.5 coverslip, 14 mm glass diameter, and Poly-D-Lysine coated glass bottoms (MatTek P35GC-1.5-14-C). Once cells adhered, they were stained with 10 μg/mL Wheat Germ Agglutinin (WGA) Alexa Fluor 555 (Thermo Fisher W32464) for 10 min at 37°C. Following staining, cells were washed twice with complete media and incubated at 37°C and 5% CO₂ for 1 hr. During this incubation, cells received the following treatments in complete media: 1 nM GSK2193874 (GSK219, MedChemExpress HY-100720) for 1 hr, 5 μM TRAM-34 (MedChemExpress HY-13519) for 1 hr, 50 μM Y-27632 dihydrochloride (ROCK Inhibitor, Sigma Aldrich Y0503) for 1 hr, or Δ74.4 mOsm/L Polyethylene glycol 300 (PEG300, Millipore Sigma 8074845000) for 15 min. For combination treatments, when multiple treatments were performed within the same dish, the first treatment was applied for its full incubation time before addition of the second treatment: order of addition was as follows: 5 μM TRAM-34 followed by Δ74.4 mOsm/L PEG300, 50 μM ROCK Inhibitor followed by Δ74.4 mOsm/L PEG300, 1 nM GSK219 followed by Δ74.4 mOsm/L PEG300, or 1 nM GSK219 followed by 5 μM TRAM-34. For combination treatments, control and single-treatment acquisitions were completed before addition of the second treatment, with each incubation time as described above. After incubation, samples were placed in an Okolab chamber at 37°C and 5% CO₂ for imaging. Time-lapse imaging was performed using confocal microscopy with a 561 nm laser for 3 hr with 1 min intervals.

### Single-Cell Tracking and Diffusivity Analysis

Cell trajectories were tracked from time-lapse movies using TrackMate (ImageJ). Single-cell centroid positions were used to compute mean squared displacement (MSD) for each trajectory. Diffusivity (*D*) was estimated by linear regression of MSD versus time delay (τ) over the first 10 lag intervals. Trajectories were included only if the short-time MSD fit yielded R²>0.8. Across two independent experiments, 87–887 cells were tracked per condition (typically >100). For each cell line and treatment, the mean single-cell diffusivity ⟨*D*⟩ was computed across included trajectories. The motility index (MI) was then calculated per cell line as the ratio of mean diffusivities under treatment versus matched control conditions (MI = ⟨D_treatment⟩/⟨D_control⟩; e.g., MI_PEG = ⟨D_PEG⟩/⟨D_control⟩ and MI_GSK219 = ⟨D_GSK219⟩/⟨D_control⟩).

### Line Profile Analysis

Immunofluorescence images of TRPV4 or pMLC, F-actin (phalloidin), and DAPI were analyzed in ImageJ. Images were concatenated, and background was subtracted from all channels. Line profiles crossing individual cells were generated to quantify protein distribution. The cortical region was defined as a ∼1.5 µm band from the cell perimeter and the nuclear region was defined by DAPI signal (region above 50% of peak nuclear intensity). The cytosol was defined as the region between the cortex and nucleus. The fraction of total cellular pMLC or F-actin signal within this cortical band was calculated for each cell. At least 12 cells were analyzed per condition across two independent experiments.

### Statistical Analysis

Statistical analyses were performed using GraphPad Prism (10.2.3), except for multiple regression analysis which was performed in MATLAB (R2022a). For correlation analysis, linear regression was performed with log₂-transformed mRNA values as the independent variable and motility index (linear scale) as the dependent variable. R^2^ (coefficient of determination) and two-sided p-values for the regression slope were calculated. To assess whether KCNN4 mRNA provided additional predictive value beyond TRPV4, multiple regression models were compared using nested F-tests: Model 1 included TRPV4 log₂ fold-change alone; Model 2 included both TRPV4 and KCNN4 log₂ fold-changes. This was performed separately for MI_PEG and MI_GSK219. For group comparisons of diffusivity and cortical enrichment measurements, two-tailed Mann-Whitney U tests were used. P<0.05 was considered statistically significant. All data are presented as mean±standard deviation (SD) unless otherwise noted. Sample sizes (n) refer to the number of cell lines analyzed for transcript-output correlations or the number of cells analyzed for imaging/motility experiments.

