## Supplementary Figures and Tables for "Log-linear scaling of TRPV4-KCNN4 transcripts tunes ROCK-dependent mechanotransduction in a DCIS progression model"

### Supplemental Information

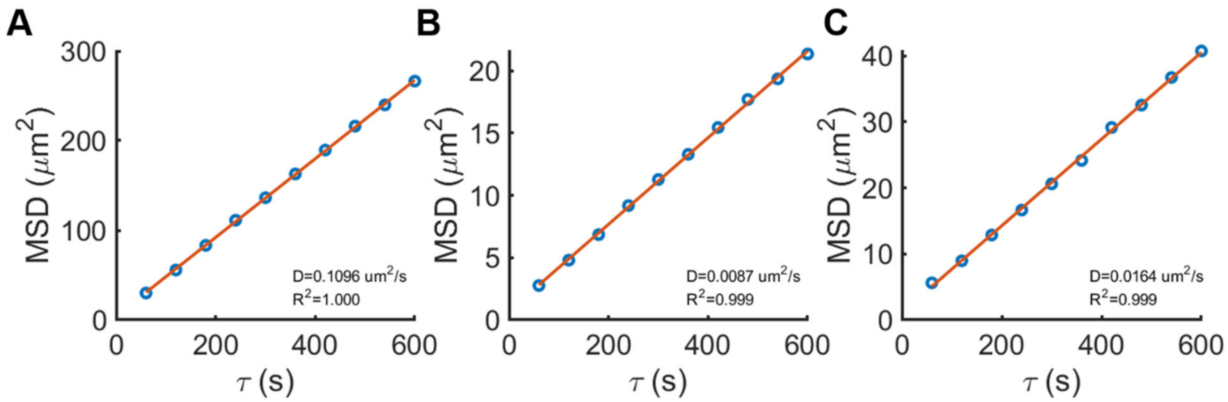


**Figure S1. Mean-squared displacement (MSD) analysis.**

Representative single-cell MSD curves from MCF10DCIS.com illustrating different diffusivity values. MSD was fit with a linear model for 2D diffusion (MSD = $4D\tau$; orange lines), where lag time $\tau = n\Delta t$ with $\Delta t = 60$ s. Diffusivity coefficients ($D$) were obtained from the fitted slopes ($D$=slope/4): 0.1096, 0.0087, and 0.0164 μm²/s. The high fit quality ($R^2 \geq 0.999$ in examples shown) supports the linear approximation. Only trajectories with fit quality $R^2 > 0.8$ were included in downstream analyses (n>100 cells per condition).

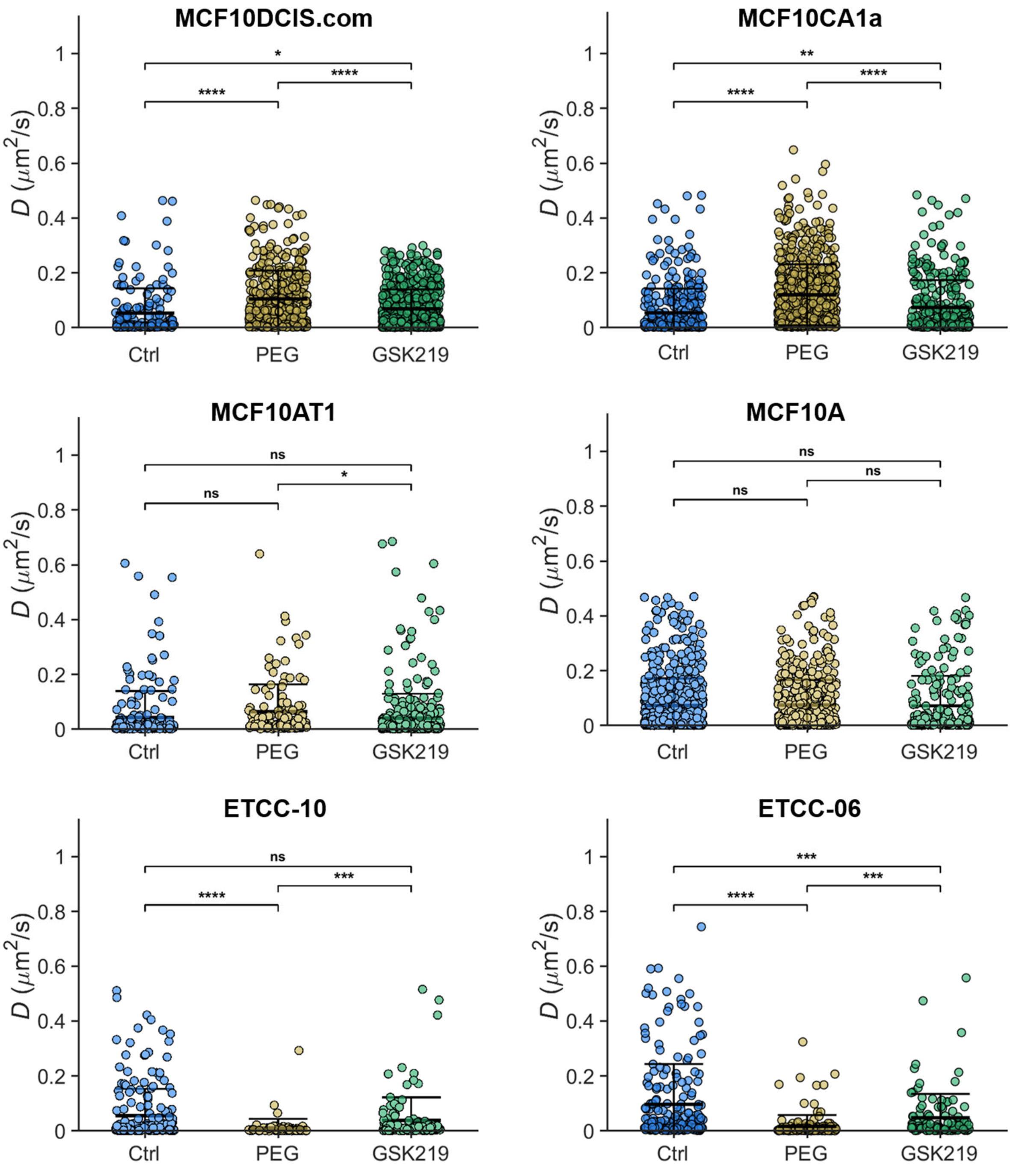


**Figure S2. Complete single-cell diffusivity distributions**

Diffusivity coefficients (*D*) for all six cell lines under control (blue), PEG 300 (PEG; gold), and GSK219 (green). Cell lines are grouped by phenotype: stress-responsive (top), none-to-weakly-responsive (middle), and negatively-responsive (bottom). Each point represents a single cell;

horizontal/vertical lines indicate mean±SD. Pairwise Mann–Whitney U tests are shown: ns, not significant; *, $p<0.05$; **, $p<0.01$; ***, $p<0.001$; ****, $p<0.0001$. $n>100$ cells per condition per cell line.

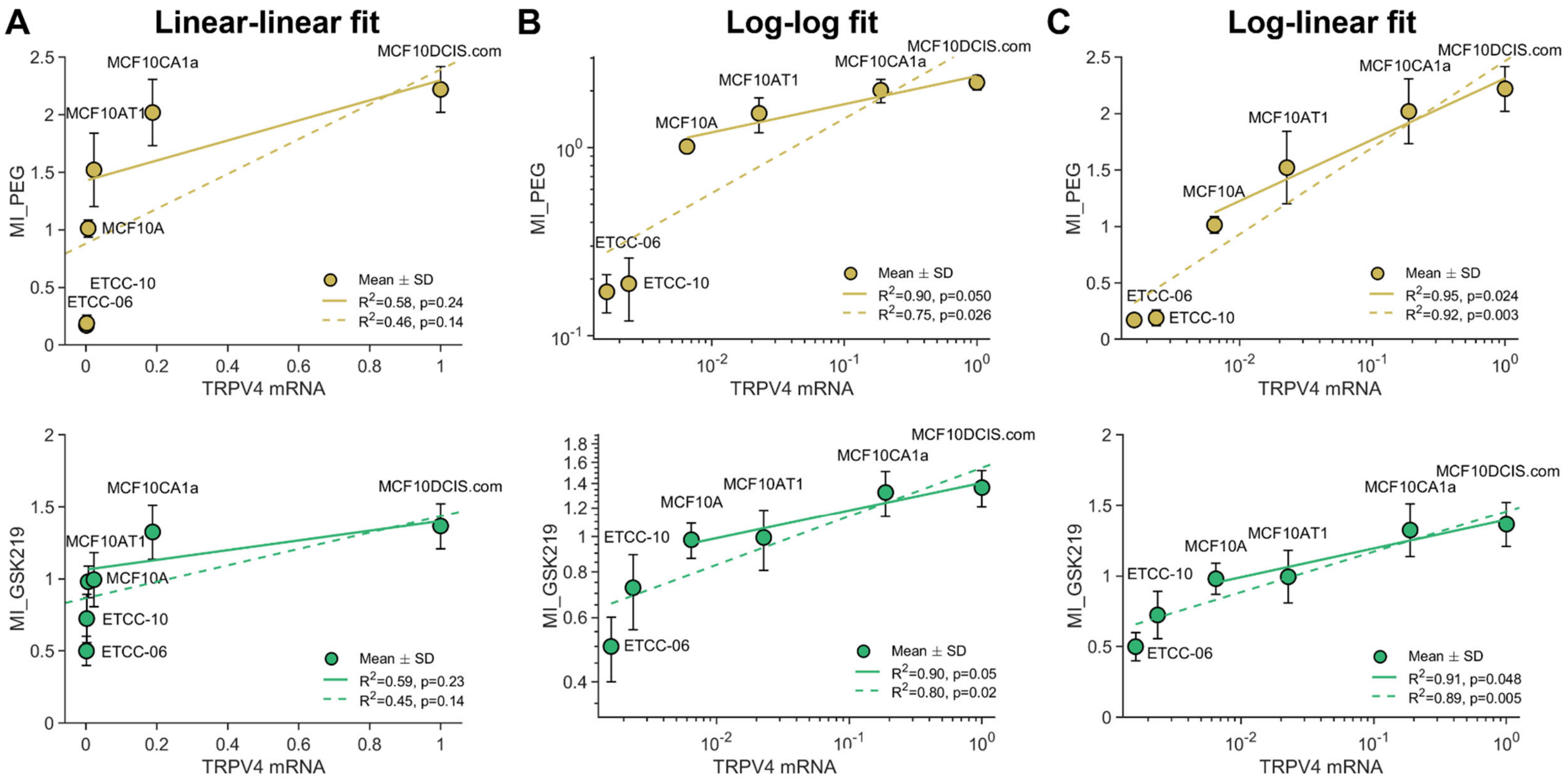


**Figure S3. Log-linear model provides superior fit for TRPV4 mRNA-motility relationships**

Comparison of alternative regression models for TRPV4 transcript abundance versus mechanotransduction output. **(A)** Linear-linear fit with both axes on linear scale. **(B)** Log-log fit with both axes on logarithmic scale. **(C)** Log-linear fit with TRPV4 mRNA on logarithmic scale and motility index (MI) on linear scale. Top row: MI_PEG (hyperosmotic stress); bottom row: MI_GSK219 (pharmacologic TRPV4 inhibition). Solid lines represent isogenic MCF10A series only (n=4 lines); dashed lines represent extended panel including patient-derived DCIS lines ETCC-06 and ETCC-10 (n=6 lines). Log-linear regression **(C)** provides superior explanatory power compared to linear-linear **(A)** or log-log **(B)** models for both triggers across both the isogenic series (PEG: $R^2$=0.95 vs 0.58 vs 0.90; GSK219: $R^2$=0.91 vs 0.59 vs 0.90) and extended panel (PEG: $R^2$=0.92 vs 0.46 vs 0.75; GSK219: $R^2$=0.89 vs 0.45 vs 0.80). Data represent mean±SD from at least three independent experiments. $R^2$ and p-values shown in each panel.

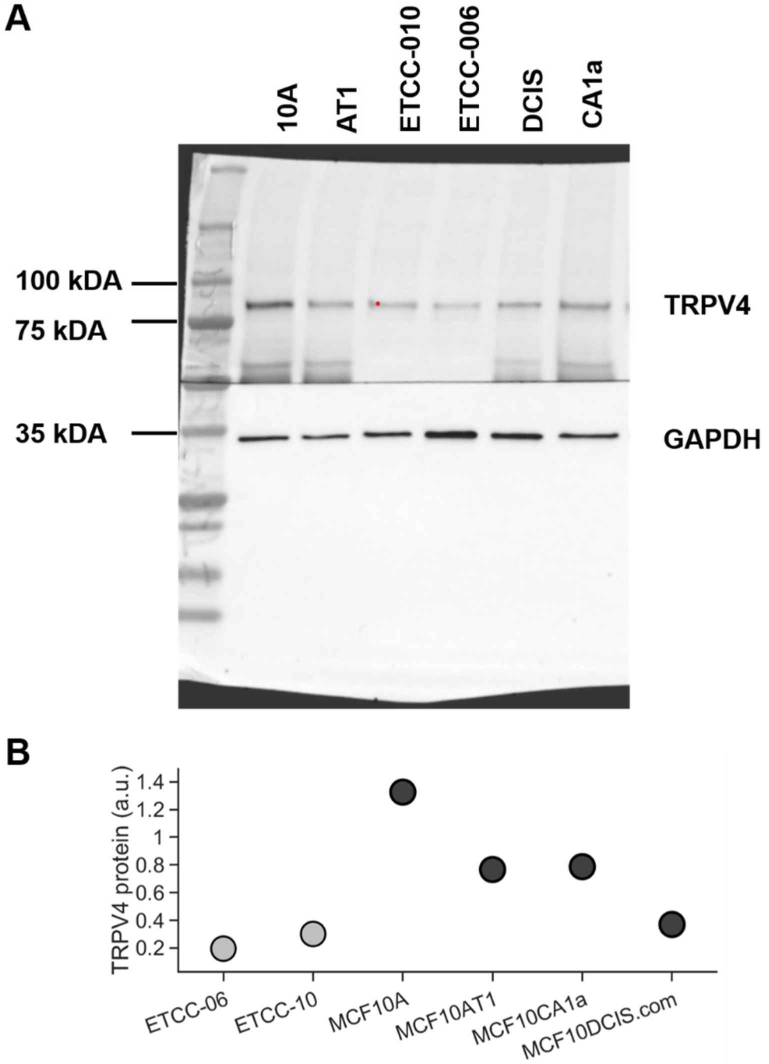


**Figure S4. Full uncropped Western blot and quantification of TRPV4 protein expression**

**(A)** Complete, uncropped WB membrane showing TRPV4 (~98 kDa; top) and GAPDH loading control (~37 kDa; bottom) across all six cell lines (MCF10A, MCF10AT1, ETCC-010, ETCC-006, MCF10DCIS, MCF10CA1a). Molecular weight ladder is shown on the left. Cells were cultured at normal density to match motility assay conditions. **(B)** Densitometric quantification of TRPV4 protein (a.u.), normalized to GAPDH, for each cell line. Cropped blot shown in **Figure 1G**.

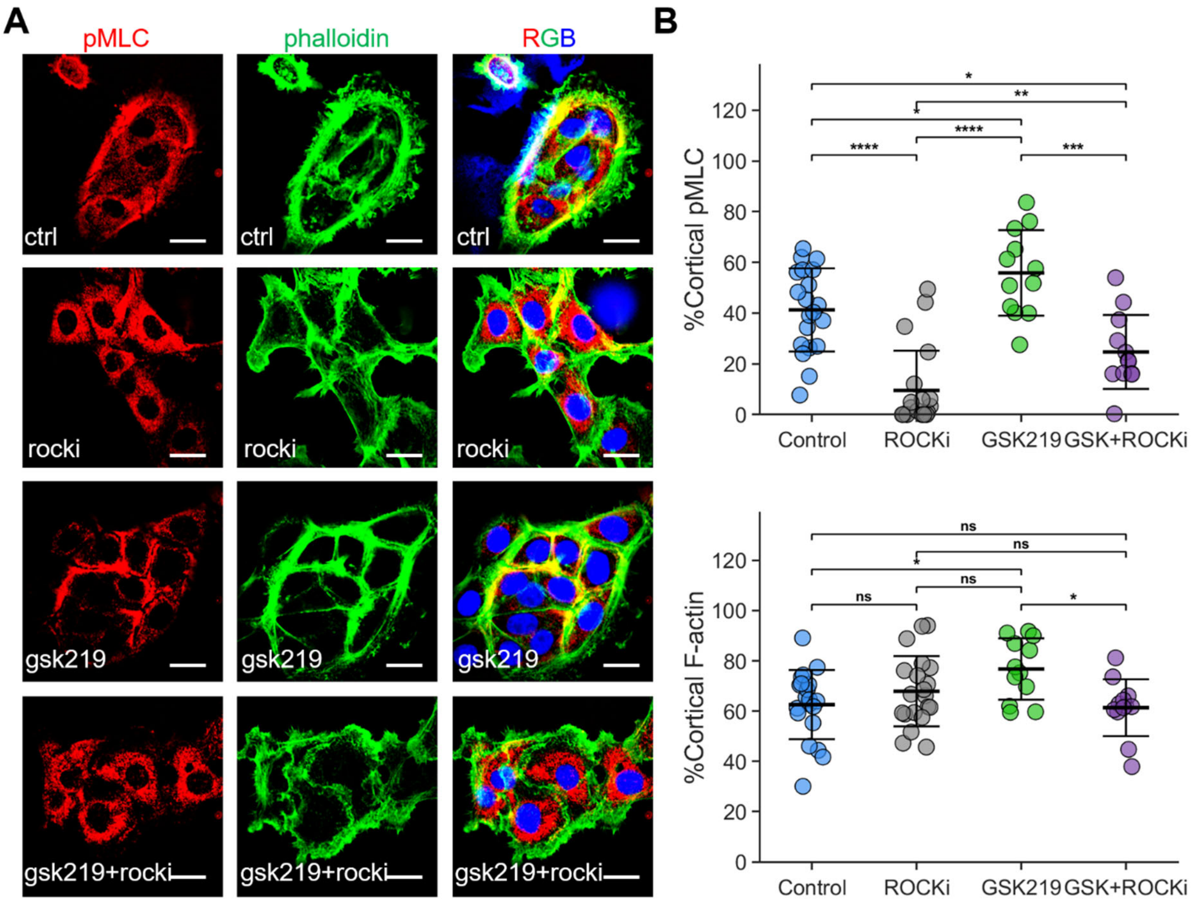


**Figure S5. TRPV4 inhibition promotes ROCK-dependent cortical contractility and actin organization.**

**(A)** Representative confocal microscopy images showing phospho-myosin light chain (pMLC, red) and F-actin (phalloidin, green) under Control, ROCK inhibition (ROCKi; Y-27632), TRPV4 inhibition (GSK219), and combined GSK219 + ROCKi conditions. Merged images include nuclei (DAPI, blue). For imaging we used 10 nM GSK219 to enhance the cortical signal; motility assays used 1 nM unless noted

**(B)** Quantification of cortical enrichment, expressed as the fraction of total cellular signal within a ~1.5 µm peripheral cortical band. Cortical pMLC2 was 41.29±16.39% in Control (n=20) and decreased with ROCKi to 9.54±15.68% (n=20). GSK219 increased cortical pMLC2 to 55.87±16.87% (n=12), and this enrichment was reduced by ROCK inhibition (GSK219+ROCKi:

24.71±14.57%, n=12). Cortical F-actin was 62.57±13.74% in Control (n=20) and 67.89±13.98% with ROCKi (n=20), while GSK219 increased cortical F-actin to 76.75±12.18% (n=12); this was reduced with ROCK inhibition (GSK219+ROCKi: 61.36±11.30%, n=12). Points represent individual cells; horizontal lines indicate mean±SD. Mann–Whitney U tests. ns p≥0.05; * p<0.05; ** p<0.01; *** p<0.001; **** p<0.0001.

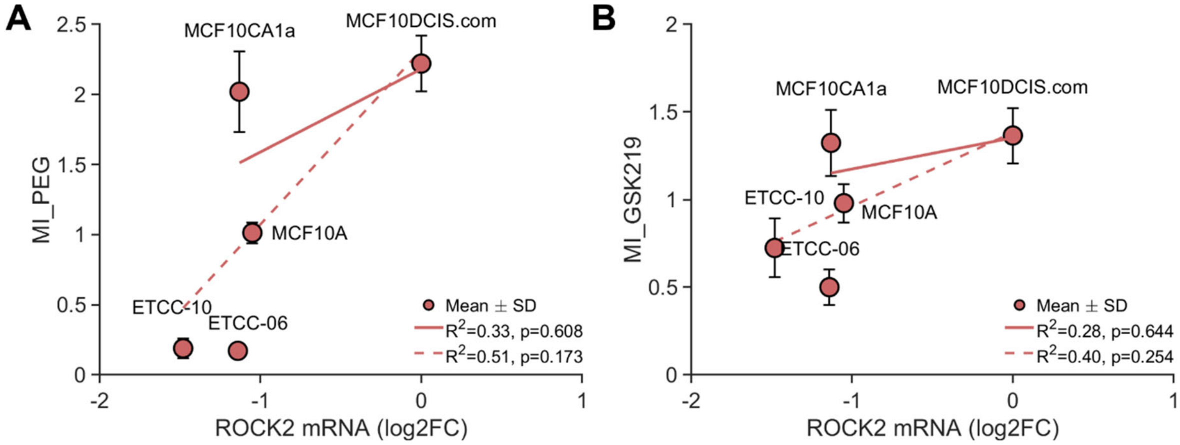


**Figure S6. ROCK2 mRNA does not correlate with mechanotransduction output despite functional requirement**

ROCK2 transcript abundance shows no relationship with mechanotransduction capacity across the breast cancer progression panel.

**(A)** Log-linear regression of ROCK2 mRNA versus MI_PEG.

**(B)** Log-linear regression of ROCK2 mRNA versus MI_GSK219. Solid lines represent isogenic MCF10A series (n=4; PEG300: $R^2$=0.33, p=0.61; GSK219: $R^2$=0.28, p=0.64); dashed lines represent extended panel including patient-derived DCIS lines (n=5; PEG300: $R^2$=0.51, p=0.17; GSK219: $R^2$=0.40, p=0.25). Despite functional requirement for ROCK activity in mediating stress-evoked cortical contractility and motility (**Figure 2**), ROCK2 mRNA levels varied minimally (2.8-fold range) and did not predict mechanotransduction output. This contrasts with plasma membrane mechanosensors TRPV4 (622-fold range, $R^2$>0.89) and KCNN4 (**Figure 1**, **Figure 3**), supporting a two-tier organization where capacity is determined by mechanosensor transcript abundance while cytoplasmic effectors are constitutively expressed and activated post-translationally. Data are mean±SD from ≥3 independent experiments. See **Supplemental Table S1** for complete gene panel.

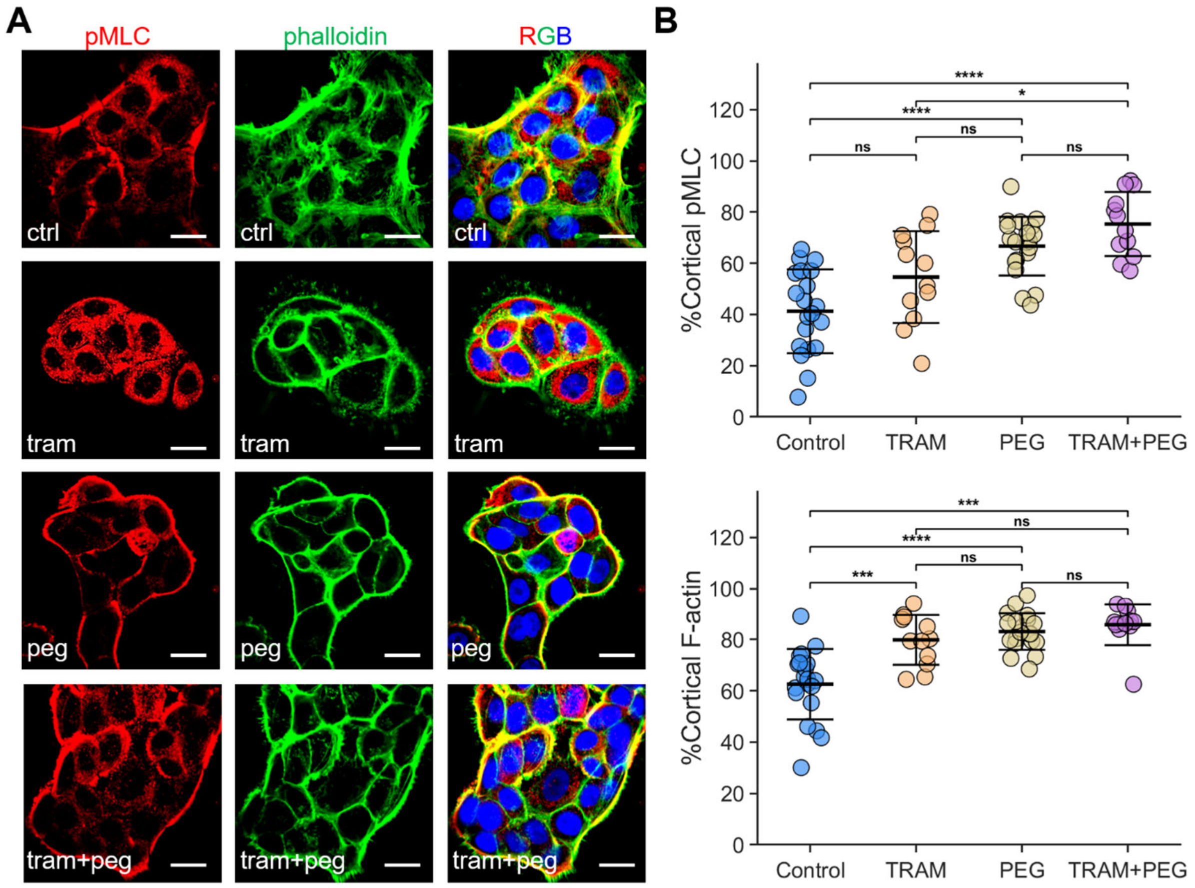


**Figure S7. TRAM-34 and hyperosmotic stress enhance cortical contractility and F-actin organization.**

**(A)** Representative confocal microscopy images showing phospho-myosin light chain 2 (pMLC, red) and F-actin (phalloidin, green) under Control, 5 μM TRAM-34 (TRAM), ∆74.4 mOsm/L PEG300 (PEG), and combined TRAM+PEG conditions. Merged images include nuclei (DAPI, blue).

**(B)** Quantification of cortical enrichment, expressed as the fraction of total cellular signal within a ~1.5 μm peripheral cortical band. Cortical pMLC2 was 41.29±16.39% in Control (n=20), 54.63±17.95% with TRAM (n=12), 66.70±11.47% with PEG (n=20), and 75.37±12.51% with TRAM+PEG (n=12). Cortical F-actin was 62.57± 13.74% in Control (n=20), 79.87±9.74% with TRAM (n=12), 83.11±7.13% with PEG (n=20), and 85.76±7.95% with TRAM+PEG (n=12). Points represent individual cells; horizontal lines indicate mean ± SD. Mann–Whitney U tests. ns, $p \geq 0.05$; * $p<0.05$; ** $p<0.01$; *** $p<0.001$; **** $p<0.0001$.

| mRNA | Subset | $R^2$ (PEG) | P (PEG) | n | $R^2$ (GSK) | P (GSK) | n |
|---|---|---|---|---|---|---|---|
| **TRPV4** | Full | 0.92 | **0.003** | 6 | 0.89 | **0.005** | 6 |
| | Iso | 0.95 | **0.024** | 4 | 0.91 | **0.048** | 4 |
| **KCNN4** | Full | 0.94 | **0.002** | 6 | 0.81 | **0.015** | 6 |
| | Iso | 0.81 | 0.099 | 4 | 0.67 | 0.184 | 4 |
| **PIEZO1** | Full | N/A | N/A | 0 | N/A | N/A | 0 |
| **PIEZO2** | Full | N/A | N/A | 0 | N/A | N/A | 0 |
| **ROCK2** | Full | 0.51 | 0.174 | 5 | 0.40 | 0.254 | 5 |
| | Iso | 0.33 | 0.608 | 3 | 0.28 | 0.644 | 3 |
| **RHOB** | Full | 0.31 | 0.329 | 5 | 0.19 | 0.464 | 5 |
| | Iso | 0.72 | 0.150 | 4 | 0.47 | 0.311 | 4 |
| **RHOC** | Full | 0.55 | 0.471 | 3 | 0.38 | 0.575 | 3 |
| **YAP1** | Full | 0.91 | 0.189 | 3 | 0.74 | 0.345 | 3 |
| **WWTR1** | Full | 0.82 | **0.033** | 5 | 0.79 | **0.045** | 5 |
| | Iso | 0.97 | 0.104 | 3 | 0.95 | 0.151 | 3 |

**Supplemental Table S1. Correlation between mechanotransduction candidate mRNA levels and mechanotransduction output across the DCIS progression panel.**

mRNA levels were quantified as log2 fold-change relative to MCF10DCIS.com and correlated with mechanotransduction output measured as motility index (MI) following either hyperosmotic stress (Δ74.4 mOsm/L PEG300; MI_PEG) or TRPV4 inhibition (GSK2193874, 1 nM; MI_GSK219). Correlations were computed by linear regression with log2 fold-change as the independent variable and MI as the dependent variable. "Iso" denotes the four-line isogenic MCF10 progression series; "Full" denotes the six-line panel (isogenic series plus two patient-derived DCIS lines). $R^2$, coefficient of determination; p, two-sided test of a nonzero regression slope; n, number of cell lines with an available log2 fold-change value for that gene in the indicated subset/condition. PIEZO1 and PIEZO2 were not present in the vendor differential-expression outputs used to extract log2 fold-change values (n=0) and were therefore not analyzed. ROCK2 log2 fold-change was unavailable for one cell line (n=5). Bold indicates significant correlation ($p<0.05$). WWTR1/TAZ exhibited an inverse association (higher expression associated with lower MI).

| Model | n | $R^2$ (PEG) | Adj. $R^2$ (PEG) | p (PEG) | $R^2$ (GSK219) | Adj. $R^2$ (GSK219) | p (GSK219) |
|---|---|---|---|---|---|---|---|
| **TRPV4** | 6 | 0.92 | 0.90 | **0.003** | 0.89 | 0.87 | **0.005** |
| **KCNN4** | 6 | 0.94 | 0.92 | **0.002** | 0.81 | 0.76 | **0.015** |
| **TRPV4+KCNN4** | 6 | 0.97 | 0.95 | **0.005** | 0.90 | 0.84 | **0.030** |
| Nested p (add KCNN4 to TRPV4) | | | | 0.099 | | | 0.64 |

**Supplemental Table S2. Multiple linear regression indicates TRPV4 and KCNN4 provide largely overlapping predictive information for mechanotransduction capacity.**

Multiple linear regression models predicting MI_PEG (hyperosmotic stress response) and MI_GSK219 (TRPV4 inhibitor response) using TRPV4 and/or KCNN4 mRNA abundance as predictors (n=6 cell lines). $R^2$ indicates proportion of variance explained; adjusted $R^2$ penalizes models for additional parameters. Model p-values reflect the overall F-test versus an intercept-only model. Nested p-values (partial F-test) evaluate whether adding KCNN4 to a model already containing TRPV4 provides significant improvement. Despite strong individual predictive power of both TRPV4 and KCNN4 for MI_PEG ($R^2 \geq 0.92$), the combined model shows only a small, non-significant increase ($\Delta R^2 \sim 0.05$, nested p=0.099). Similarly, for MI_GSK219, adding KCNN4 to TRPV4 provides no significant improvement ($\Delta R^2 \sim 0.01$, nested p=0.64). This lack of additive predictive value is consistent with tight transcriptional co-regulation ($R^2 = 0.83$, **Figure 3E**), supporting the interpretation that TRPV4 and KCNN4 function as a coordinated mechanosensor module rather than independent predictors.
